# A multi-omics approach to identify the impact of miR-411ed on NSCLC TKI resistance

**DOI:** 10.64898/2026.03.31.715663

**Authors:** Daniel Del Valle Morales, Giulia Romano, Michela Saviana, Patrick Nana-Sinkam, Giovanni Nigita, Mario Acunzo

**Author notes:** Corresponding authors: Giovanni Nigita; Mario Acunzo.

## Abstract

Tyrosine Kinase inhibitors (TKIs) are widely used as effective chemotherapeutic agents for treating patients with EGFR-mutated NSCLC. Unfortunately, after treatment, patients eventually develop resistance to TKI therapy. The most common resistance mechanism for the TKI Osimertinib is the overexpression of the MET Proto-Oncogene, Receptor Tyrosine Kinase (MET). We previously demonstrated that miR-411-5p A-to-I edited at position 5 (miR-411ed) can directly target MET in A549 and H1299 cells. MiR-411ed in combination with Osimertinib reduced cell proliferation in two TKI resistant EGFR-mutated cell lines: HCC827R and PC9R. MiR-411ed did not downregulate MET expression in HCC827R, suggesting an alternative mechanism for TKI response. In this study, we aim to identify the mechanism of miR-411ed TKI response using a multi-omics approach of RNAseq and protein mass spectrometry. In our cellular model, we identified miR-411ed affected genes independent of MET activity, resulting in 211 genes (RNAseq) and 36 proteins (proteomics). Pathway analysis identified an increase in interferon signaling for RNAseq and combined omics, and a decrease in ERK/MAPK signaling in proteomics. Using the IsoTar target prediction tool, we identified STAT3 as a key regulator and confirmed STAT3 protein downregulation upon transfection with miR-411ed. We further investigated the effect of miR-411ed in vivo, observing a reduction in tumor size with miR-411ed in combination with Osimertinib but not with miR-411ed or Osimertinib treatment alone, confirming the effectiveness of miR-411ed in TKI response.

## Introduction

Epidermal Growth Factor Receptor (EGFR) mutation is a key driver of Non-Small Cell Lung Cancer (NSCLC) progression, with a prevalence of 10-20% in Western populations and an increased prevalence (50%) in Asian populations^1^. Osimertinib is a third-generation tyrosine kinase inhibitor (TKI) used to treat patients with EGFR-positive lung cancers. Despite initial response, patients eventually develop resistance to Osimertinib. The most common mechanism of resistance is the overexpression of Proto-Oncogene Proteins c-met (MET), which accounts for 20% of cases^2^. MET overexpression bypasses EGFR TKIs by reactivating the ERK/MAPK and AKT pathways.

MicroRNAs (miRNA/miR) have been shown to play a major role in the development of TKI resistance in lung cancer^3^. MiRNAs are short regulatory RNAs that decrease gene expression by the direct binding of the miRNA “seed region” to the 3’UTR of their mRNA target, inhibiting its translation or promoting RNA degradation. Additionally, miRNAs can undergo posttranscriptional modifications such as A-to-I editing that alter seed region base pairing and their function^4^. The role of miRNA editing in the pathogenesis of lung cancer has yet to be fully elucidated.

Previously, we identified hypoediting on position 5 of the seed region of miR-411-5p (miR-411ed), which was reduced in NSCLC patients^5^. We demonstrated that gain-of-function of miR-411ed reduced MET expression in NSCLC cell lines A549 and H1299 compared to wt miR-411-5p^4^. Additionally, miR-411ed, in combination with Osimertinib, reduced proliferation in two TKI resistant cell lines: PC9R (MET-independent) and HCC827R (MET-amplified). Despite demonstrating direct targeting of MET, miR-411ed does not reduce MET expression in resistant cells. Interestingly, miR-411ed reduces p-ERK in resistant cells, suggesting an alternative mechanism for inducing a TKI response.

In this short communication, we used a multi-omics approach to identify miR-411ed dysregulated genes independent of MET activity and examined the effectiveness of miR-411ed in combination with Osimertinib *in vivo*.

## Material and Methods

### Cell culture

HCC827R cells were grown and transfected as in Romano et al.^4^. The MET inhibitor PF04217903 (5µM) or DMSO was added to transfected cells overnight before cell harvest. Stable HCC827R cells used for *in vivo* experiments were generated by stably transfecting the cells with either 1 µg of pGreenPuro (EF1α) shRNA Lentivector (SI506A-1; SBI) as a control (Ctr) or a custom miR411-targeting pGreenPuro (EF1α) shRNA Lentivector (SI506A-1; SBI) plasmid. Transfections were performed using Lipofectamine 3000 (ThermoFisher # A12621). 48hrs post-transfection, cells were treated with puromycin (0.75µg/mL) and maintained under selection for the duration of all experiments.

### Western Blot and RNA extraction

Protein and RNA extraction were performed as described in Romano et al. The antibodies used are P-MET (Invitrogen #700139), MET (Cell Signaling #4560), STAT3 (Cell Signaling #7907), and GAPDH (Cell Signaling #3683).

### RNAseq

Sequencing libraries were prepared from the extracted RNA using the Illumina Stranded mRNA Prep library kit following standard protocols. Library concentration and quality were assessed by absolute quantification using QIAGEN QIAcuity digital PCR. Equimolarly pooled libraries were sequenced on the Illumina NextSeq 2000 platform using the recommended Illumina P3-200 cycles sequencing chemistry and protocols to generate high-quality sequencing data suitable for downstream analysis.

### LC-MS/MS

The samples were digested using the Preomics iST sample clean-up protocol. To the sample containing approximately 100µg of protein, 50µl of lysis buffer was added and mixed, followed by an incubation for 10 minutes at 95ºC at 1000rpm and sheared using a sonicator (4 cycles; sec ON/OFF). Samples were spun and transferred to the cartridge provided in the Kit. 50µl of DIGEST solution was added to the mixture, which was incubated at 37°C for 3hrs at 500 rpm. After digestion, 100µl of STOP solution was added and mixed. The digest was centrifuged at 3800rcf; 3min and washed with 200µl of WASH 1 and 200ul of WASH 2 solution followed by centrifugation after each wash. The cartridge was placed into the fresh collection tube, and 100ul of ELUTE solution was added, then centrifuged at 3800rcf for 3min, and then repeated. The elutes were placed in a vacuum evaporator at 450 C until completely dried. The samples were further cleaned using the Phoenix peptide clean-up kit. LC-MS/MS analysis was performed using a 480 Exploris tandem mass spectrometer (Thermo) coupled to a Neo Vanquish nanoflow UPLC system (Thermo). The LC-MS/MS system was fitted with an EasySpray ion source and an Acclaim PepMap 300µm x 5mm nanoviper C18 5µm x 100Å pre-column in series with an Acclaim PepMap Neo 75µm x 500mm C18 2µm bead size (Thermo). The mobile phase consists of Buffer A (0.1% formic acid in water) and Buffer B (80% acetonitrile in water, 0.1% formic acid). 500ng of peptides were injected onto the above column assembly and eluted with an acetonitrile/0.1% formic acid gradient at a flow rate of 350 nL/min over 2 hours. The nano-spray ion source was operated at 1.9 kV. The digests were analyzed using a data-dependent acquisition (DDA) method, acquiring a full scan mass spectrum (375-1500m/z) at an AGC = 300% and an ITmax = Auto followed by 10 HCD tandem mass spectra per second from 78-6000 m/z at an NCE = 30%, AGC = Standard, and ITmax = Auto.

### In vivo Tumor Studies

Animal experiments were conducted in accordance with a protocol approved by Virginia Commonwealth University Institutional Animal Care and Use Committee. NSG (NOD.Cg-Prkdcscid Il2rgtm1Wjl/SzJ) mice, 6 weeks of age, were obtained from VCU Cancer Mouse Models Core, which maintains colonies originating from founders purchased from The Jackson Laboratory. All cell lines used for implantation were confirmed to be free of Corynebacterium bovis and mycoplasma contamination before injection.

Mice were injected subcutaneously in the right flank with 2.5 × 10^6^ cells. Cells were resuspended in 100 μL of a 1:1 mixture of PBS and basement membrane extract (Bio-Techne). Body weight and tumor dimensions were recorded twice weekly. Tumor volume (V) was calculated using the standard formula: V (mm^3^) = 0.5 × length × (width)^2^.

Once tumors reached approximately 50 mm^3^, a total of 20 mice per cell line (10 males and 10 females) were randomized into four study groups (n = 10 per group) based on body weight and tumor volume using Studylog software. Mice then received oral gavage with either vehicle control or Osimertinib (25 mg/kg) three times weekly. Dose volume was adjusted according to individual body weight at 8 µL/g. The treatment period lasted 45 days (study Day 19 to Day 63).

The two vehicle groups were euthanized on Day 47 due to tumor burden reaching ∼2000 mm^3^. The remaining Osimertinib-treated groups were euthanized at the planned study endpoint on Day 63, at which point tumor volumes ranged from ∼100–500 mm^3^. At euthanasia, mice were imaged, and primary tumors were excised, weighed, and processed. Tumors were divided such that half were fixed in formalin for histologic analysis, and half were flash-frozen for molecular studies.

### Immunohistochemistry

Immunohistochemical staining (IHC) was performed by the VCU Tissue and Data Acquisition and Analysis Core using the Leica Bond RX autostainer. Heat-induced epitope retrieval was carried out for 20 minutes with either BOND solution 2 (EDTA-based, pH 9.0) for Ki-67 and Cleaved Caspase-3 or BOND solution 1 (citrate-based, pH 6.0) for Transferrin Receptor. Ki-67 was diluted in Bond Primary Antibody Diluent (Leica Biosystems, AR9352), while Cleaved Caspase-3 and Transferrin Receptor were diluted in PowerVision Universal IHC Blocking Diluent (Leica Biosystems, PV6123). Blocking for Transferrin Receptor and Cleaved Caspase-3 was performed for 20 minutes using the same PowerVision diluent.

Primary antibody incubation times were as follows: Ki-67 for 15 minutes, Transferrin Receptor for 30 minutes, and Cleaved Caspase-3 for 45 minutes. Antibodies used were: Ki-67 (1:3000, Cell Signaling Technology #9027), Cleaved Caspase-3 (1:500, Cell Signaling Technology #9664), and Transferrin Receptor (1:6000, ThermoFisher #13-6800). After primary antibody incubation, slides were treated with secondary antibodies for 8 minutes, followed by HRP detection using DAB for 10 minutes with the BOND Polymer Refine Detection kit (Leica Biosystems, DS9800). Stained slides were imaged using the PhenoImager HT (Akoya Biosciences).

### Data analysis

RNAseq data were processed through a standard upstream and downstream analysis workflow. Briefly, adapter trimming and quality control were performed using Trim Galore (v0.6.6)^6^, which incorporates cutadapt (v3.5)^7^. The resulting high-quality reads were aligned to the human reference genome hg38 using STAR (v2.7.10a)^8^, with transcript annotation based on the GENCODE primary assembly annotation (v44). Transcript-level quantification was then carried out using featureCounts (v2)^9^ with the same GENCODE v44 primary assembly annotation. Lowly expressed transcripts were filtered out, and only transcripts with a geometric mean expression of ≥1 RPKM were retained for downstream analyses. Raw counts from the filtered transcripts were normalized using the trimmed mean of M-values (TMM) method, and differential expression analysis was performed with the limma R package (v3)^10^, using the eBayes function to identify transcripts with significant differential expression. The LC-MS/MS were analyzed in Proteome Discoverer (ver 3.0) using the Sequest HT search algorithm and the UniProt Human protein database. Proteins were identified with an FDR <0.01, and quantification was based on peptide intensities. Raw protein abundances were normalized in Proteome Discoverer using the “Total Peptide Abundance” method. Normalized abundances were filtered for number of peptides ≤ 2, number of unique peptides ≤ 1, and protein FDR confidence high for all groups. Genes with missing values were filtered out from the analysis. The filtered normalized abundances were analyzed for differential expression using the R package promor^11^. Transcripts without annotated gene IDs were removed from downstream analysis. Gene enrichment analysis was performed using Ingenuity^®^ Pathway Analysis (IPA^®^) (v01-23-01) software on the DEGs for both RNAseq and proteomics. Prediction of miR-411ed targets was performed using isoTar^12^ v1.2.1 using all 5 databases (PITA, RNAhybrid, TargetScan, miRanda, miRmap) and a consensus of at least one database for each predicted target.

## Results

We employed a multi-omics approach to identify genes altered with miR-411ed overexpression. HCC827R cells were transfected with miR-411ed and treated with either DMSO or the MET inhibitor to compare and identify the MET-independent miR-411ed genes. The cell lysates were analyzed using next-generation sequencing (NGS) and mass spectrometry (proteomics) (Fig 1A). Differentially expressed genes (DEG, <0.05 p.adj. RNaseq [Fig 1B] /p.value proteomics [Fig 1C] and log2FC < +/-0.58) from the comparison miR-411ed vs. scr DMSO and miR-411ed vs. scr MET-inhibited were intersected to identify MET-independent miR-411ed DEGs from RNAseq (Fig. 1B, Table S1) and proteomics (Fig. 1C, Table S2). Genes with annotated gene IDs were considered for the intersection and downstream analysis. The MET-independent DEGs were subjected to canonical pathway analysis using Ingenuity Pathway Analysis (IPA, QIAGEN Inc., https://digitalinsights.qiagen.com/IPA). The analysis of the RNA (Fig. 1D, Table S3) identified an increase in interferon signaling pathways. The analysis of the proteins identified the ERK/MAPK signaling pathway as the most significantly affected and downregulated pathway (Fig. 1E, Table S3).

**Figure 1.**
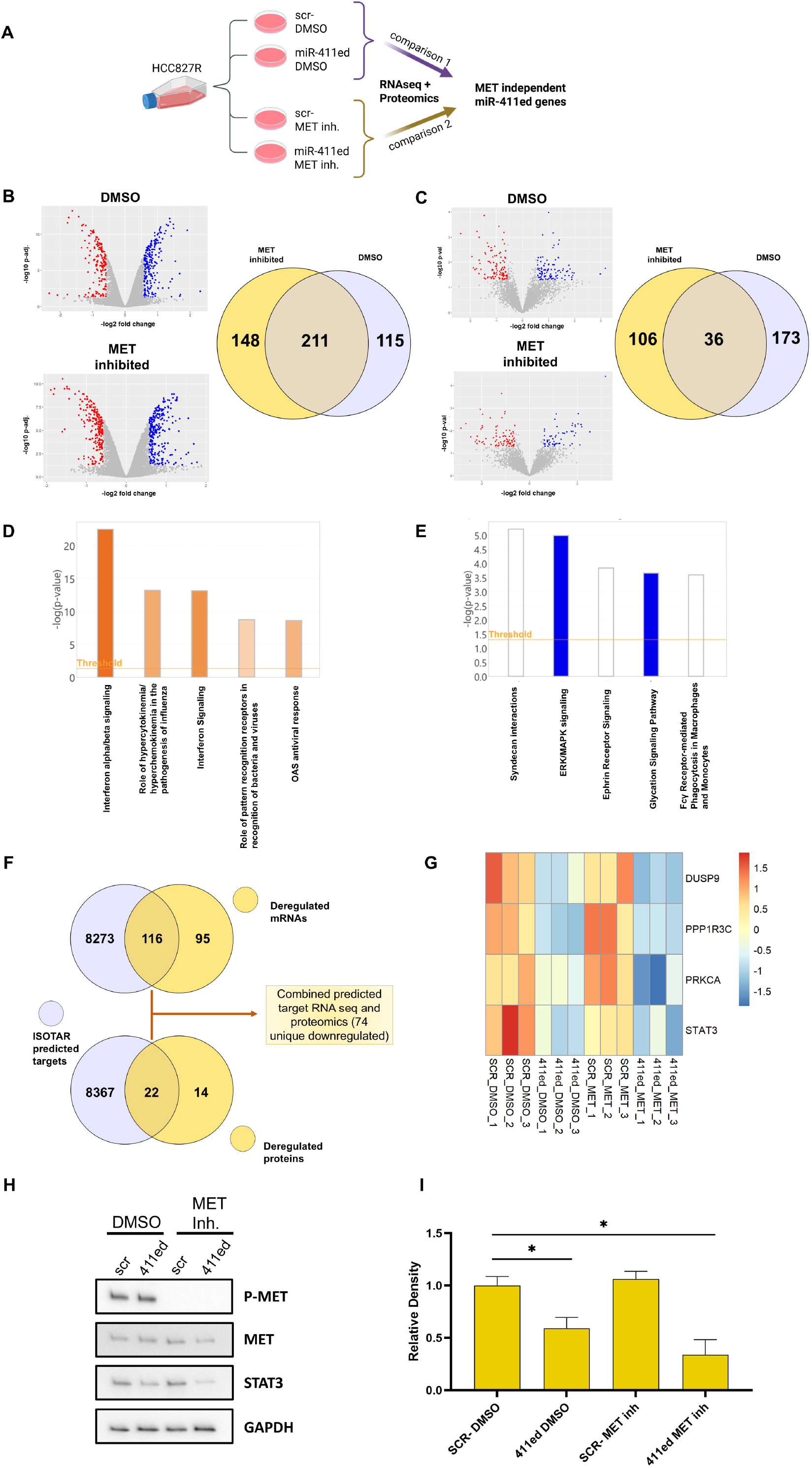
Multi-omic approach to identify miR-411ed MET-independent DEGs. A) Schema for the multi-omics approach scr-/miR-411ed DMSO (comparison 1) and scr-/miR-411ed MET-inhibited (comparison 2). B) Volcano plot of RNAseq DMSO and MET-inhibited scr vs. miR-411ed (left). Significantly decreased genes (<0.05 p.adj. and log2FC < -0.58) are in red and significantly increased genes (<0.05 p.adj. and log2FC > 0.58) are in blue. Venn diagram comparison of MET-inhibited and DMSO DEGs (right). C) Volcano plot of proteomics DMSO and MET-inhibited scr vs. miR-411ed (left). Significantly decreasing genes (<0.05 p.val. and log2FC < -0.58) are in red and significantly increasing genes (<0.05 p.val. and log2FC > 0.58) are in blue. Venn diagram comparison of MET-inhibited and DMSO DEGs (right). D) IPA analysis of the top 5 pathways for RNAseq MET-independent genes. E) IPA analysis of the top 5 pathways for proteomics MET-independent genes. F) IsoTar prediction of miR-411ed targets overlapped with RNAseq and proteomics DEGs. G) Heatmap of miR-411ed targets identified in the ERK/MAPK pathway scaled to z-score. H) Western blot of MET, P-MET, STAT3 and GAPDH as loading control. I) Relative densitometry normalized to GAPDH of STAT3. Student’s T-test determined the P-value. *= p.val< 0.05.

To identify the miR-411ed targets in the MET-independent DEGs, we employed the isoTar tool^12^. The predicted targets were filtered down to unique gene IDs and compared with the RNAseq and proteomic DEGs (Fig. 1F, Table S4), yielding 74 downregulated DEGs predicted to be targets of miR-411ed. With the identification of the ERK/MAPK signaling pathway in the proteomics analysis and our previous observation of downregulated p-ERK following miR-411ed transfection, we compared the miR-411ed-predicted targets to this pathway. We identified PRKCA, DUSP9, PPP1R3C, and STAT3 as predicted miR-411ed targets in this pathway (Fig. 1G, Table S4). STAT3 has been shown to mediate TKI resistance, and its downregulation has been associated with enhanced TKI sensitivity in NSCLC^13^. The western blot on HCC827R scr/miR-411ed and DMSO/MET inhibitor showed downregulation of STAT3 only when miR-411ed is transfected (Fig. 1H and 1I, Fig. S1), confirming the predicted downregulation of STAT3 with miR-411ed.

We evaluated the effectiveness of miR-411ed overexpression in combination with Osimertinib *in vivo* using a xenograft mouse model derived from HCC827R cells. HCC827R was stably transfected with an empty plasmid or one overexpressing miR-411ed and injected into the flank of nude mice treated with vehicle or Osimertinib (Fig. 2A). The vehicle group of mice showed no difference in tumor growth between the scr and miR-411ed mice (Fig. 2B). In the group treated with Osimertinib, we observed a reduction in tumor size of the miR-411ed mice compared to scr (Fig. 2C), confirming the effectiveness of the combination treatment *in vivo*. To test the mechanism of tumor reduction, we performed immunohistology staining for the proliferation marker Ki-67 and apoptosis. We did not observe a significant difference in Ki-67 signal between Osimertinib scr- and miR-411ed groups. When mice were stratified by sex, a decreasing trend was observed in miR-411ed male mice but not in female mice. No change in apoptosis was observed (Fig. S2).

**Figure 2.**
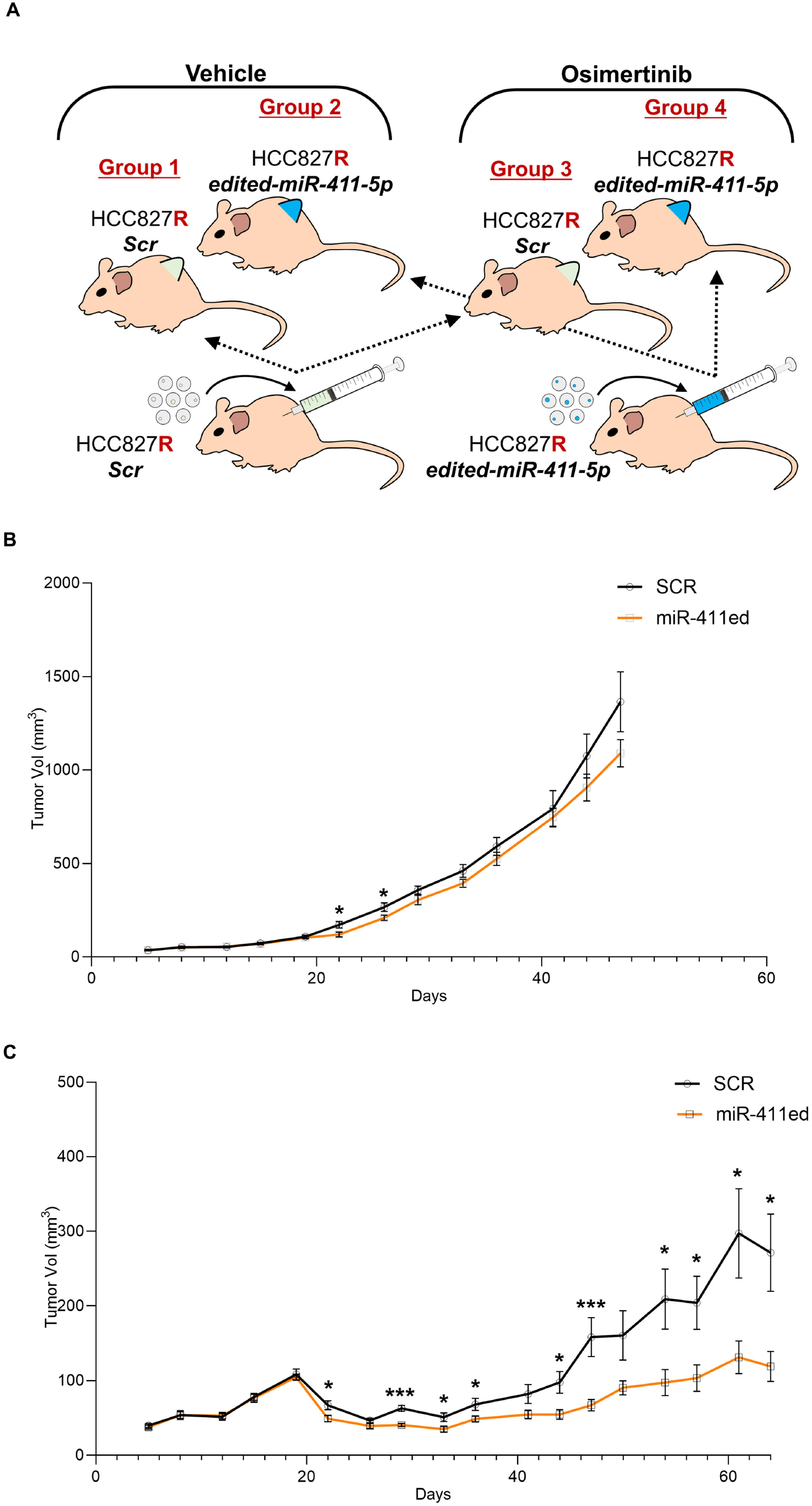
*In vivo* treatment of miR-411ed in combination with Osimertinib. A) Schema of the four treatment groups: group 1: scr HCC827R vehicle, group 2: miR-411ed HCC827R vehicle, group 3: scr HCC827R Osimertinib, group 4: miR-411ed HCC827R Osimertinib. B) Tumor volume of Group 1 and Group 2 (vehicle-treated). Mice were sacrificed on day 41 due to tumor burden. P-value was determined by Student’s t-test *= p.val< 0.05. C) Tumor volume of Group 3 and Group 4 (Osimertinib treated). Mice were treated with Osimertinib on day 19. P-value was determined by Student’s t-test *= p.val< 0.05. ***= p.val< 0.001.

## Discussion

TKI inhibitors, such as Osimertinib, are the standard of treatment for NSCLC patients harboring EGFR mutations. Despite initial response, patients develop drug resistance with MET overexpression, accounting for 20% of Osimertinib resistance cases. Dysregulation of miRNAs is one of the mechanisms for MET overexpression^2^, and A-to-I editing of miRNAs adds a new layer of regulation of MET overexpression.

We previously identified hypo-editing of miR-411ed in NSCLC tumors and in circulation^5^, demonstrated that miR-411ed directly targets MET in NSCLC cell lines^4^, and miR-411ed restores TKI sensitivity in MET-amplified (HCC827R) and MET-independent (PC9R) TKI resistant cell lines^4^. Here, we performed a multiomic analysis of RNAseq and proteomics to capture the full scope of miR-411ed DEGs and compared the DEGs -/+ MET inhibitor to identify MET-independent DEGs (Fig. 1A-C). Pathway analysis identified an activation of interferon alpha signaling in RNAseq (Fig. 1D) and inhibition of the ERK/MAPK pathway in proteomics (Fig. 1E). Reactivation of the ERK signaling pathway is common across the various mechanisms of TKI resistance^14^ and, consistently with our previous observation that miR-411ed overexpression reduces p-ERK in HCC827R and PC9R cells^4^, the MET-independent DEGs showed a decrease in ERK/MAPK signaling in the proteomics.

After combining the isoTar prediction for miR-411ed with both the RNAseq and proteomics DEGs, we identified 74 downregulated predicted targets (Fig. 1F). We selected predicted miR-411ed targets in the ERK/MAPK signaling pathway and identified PRKCA, DUSP9, PPP1R3C, and STAT3 as miR-411ed targets (Fig. 1G) and confirmed STAT3 protein downregulation after miR-411ed transfection (Fig. 1H-I). The downregulation of STAT3 by miR-411ed inhibits a key survival pathway for various TKI resistance mechanisms as STAT3 is involved in tumorigenesis, proliferation, and TKI drug resistance in NSCLC by upregulating survival genes such as c-Myc, cyclin D1, and Bcl-XL^13^. Additionally, STAT3 is an inhibitor for interferon alpha^13^, which is consistent with our RNAseq pathway results. Further investigations are needed to better understand the mechanism by which STAT3 contributes to miR-411ed-induced Osimertinib sensitivity in NSCLC.

Notably in our analysis, STAT3 was absent from the MET-independent RNAseq and was identified only by our proteomic analysis. The available miRNA target databases primarily focus on canonical miRNAs, limiting the study of edited miRNAs to target predictions and transcriptomic validation. This result indicates that the traditional approach of using RNAseq alone for miRNA target identification is insufficient to fully characterize miRNA targets. Thus, multi-omics analysis captured the full range of DEGs affected by miR-411ed and is an ideal approach to characterize the miRNA targets of edited miRNAs.

When treated in combination with Osimertinib, miR-411ed induced a TKI response in resistant NSCLC cells *in vivo*. We found no differences in tumor growth between scr- and miR-411-ed mice in the untreated group, consistent with our observation *in vitro*^4^. Conversely, when the mice were treated with Osimertinib, we observed reduced tumor growth in the miR-411ed mice compared to the scr mice. This result demonstrates that miR-411ed, in combination with Osimertinib, reduces tumor growth, paving the way for the potential application of miRNA as an adjuvant therapeutic strategy in TKI resistant lung cancer. We did not observe any significant differences in tumor proliferation or apoptosis, which may be limited to the number of mice tested. A more extensive *in vivo* study with a larger sample set would aid in elucidating the mechanism for tumor reduction and investigating whether STAT3 may play a role in the *in vivo* response.

## Conclusion

In summary, we identified MET-independent miR-411ed dysregulated genes in MET-amplified HCC827R TKI resistant cells. We confirmed the downregulation of a key miR-411ed predicted target STAT3, which was only identified through our multiomic approach highlighting the advantage of our approach in characterizing edited miRNA DEGs. Additionally, we demonstrated the effectiveness of miR-411ed treatment in combination with Osimertinib *in vivo*.

## Acknowledgements

The proteomic study for this research project was generated by the VCU Massey Comprehensive Cancer Center Proteomics Shared Resource, supported, in part, with funding from NIH-NCI Cancer Center Support Grant P30 CA016059. The RNAseq included in this study was generated at the Genomics Core facility at Virginia Commonwealth University. Services and products in support of the research project were generated by the Virginia Commonwealth University Cancer Mouse Models Core Laboratory and the Tissue and Data Acquisition and Analysis Core Laboratory, supported, in part, with funding to the Massey Cancer Center from NIH-NCI Cancer Center Support Grant P30 CA016059. Dr. Nigita was supported by the NIH-NCI (R21CA287180) for this project.

## CRediT authorship contribution statement

**Daniel del Valle Morales:** Original draft preparation, Investigation, Validation, and Visualization. **Giulia Romano:** Project administration, Investigation, Methodology, Writing-Review and Editing. **Michela Saviana**: Writing-Review and Editing, Investigation. **Patrick Nana-Sinkam**: Writing-Review and Editing, Resources, and Funding acquisition. **Giovanni Nigita**: Software, Data curation, Writing-Reviewing and Editing, Formal analysis, Recourses, Funding acquisition. **Mario Acunzo:** Writing-Reviewing and Editing, Supervision, Conceptualization, Resources, and Funding acquisition.

## Data Statement

The data will be provided upon request to the authors

## Supplemental figures

Table S1 NGS analysis

Table S2 Proteomic analysis

Table S3 IPA results

Table S4 IsoTar prediction miR-411ed

Figure S1 Full western blot of STAT3 related to figure 1H-1I

Figure S2 Immunohistology on proliferation marker Ki-67. A) Ki-67 intensities in scr-Osimertinib and miR-411ed Osimertinib mice groups (n=10). B) Ki-67 intensities separated based on mouse sex.

