## Supplementary figures and images for "A multi-omics approach to identify the impact of miR-411ed on NSCLC TKI resistance"

### Supplemental Figure 1

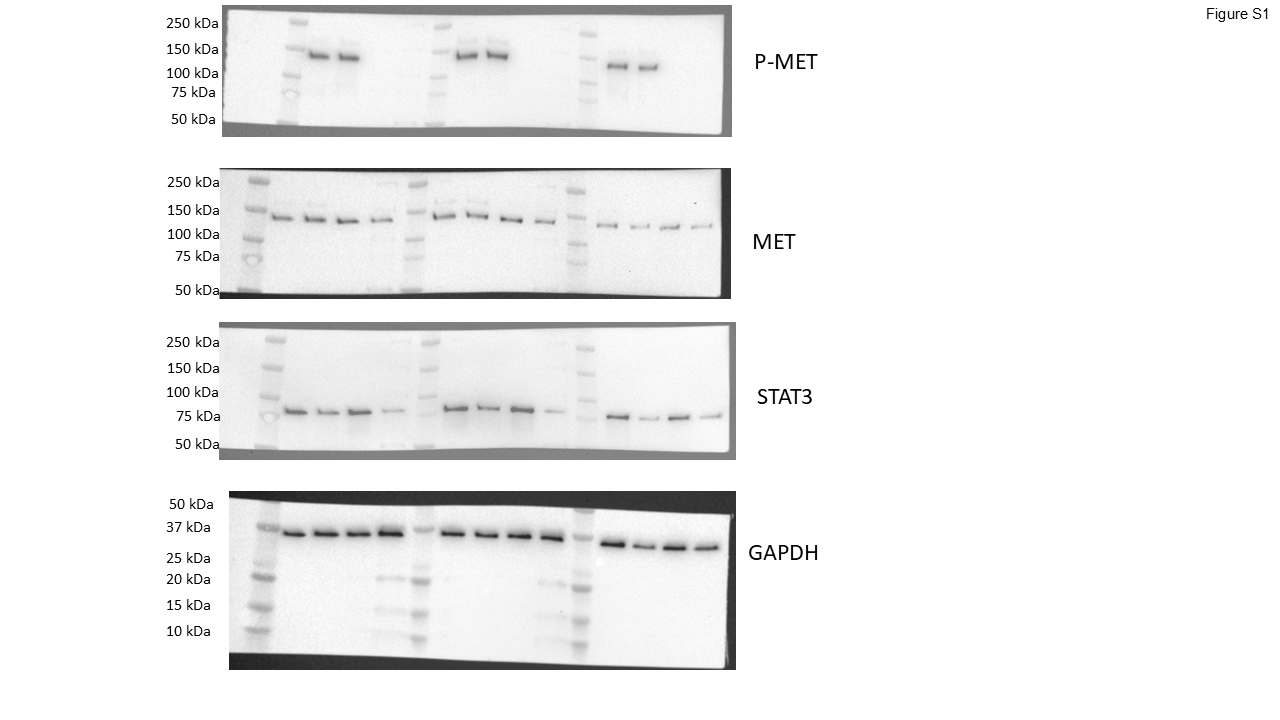

### Supplemental Figure 2

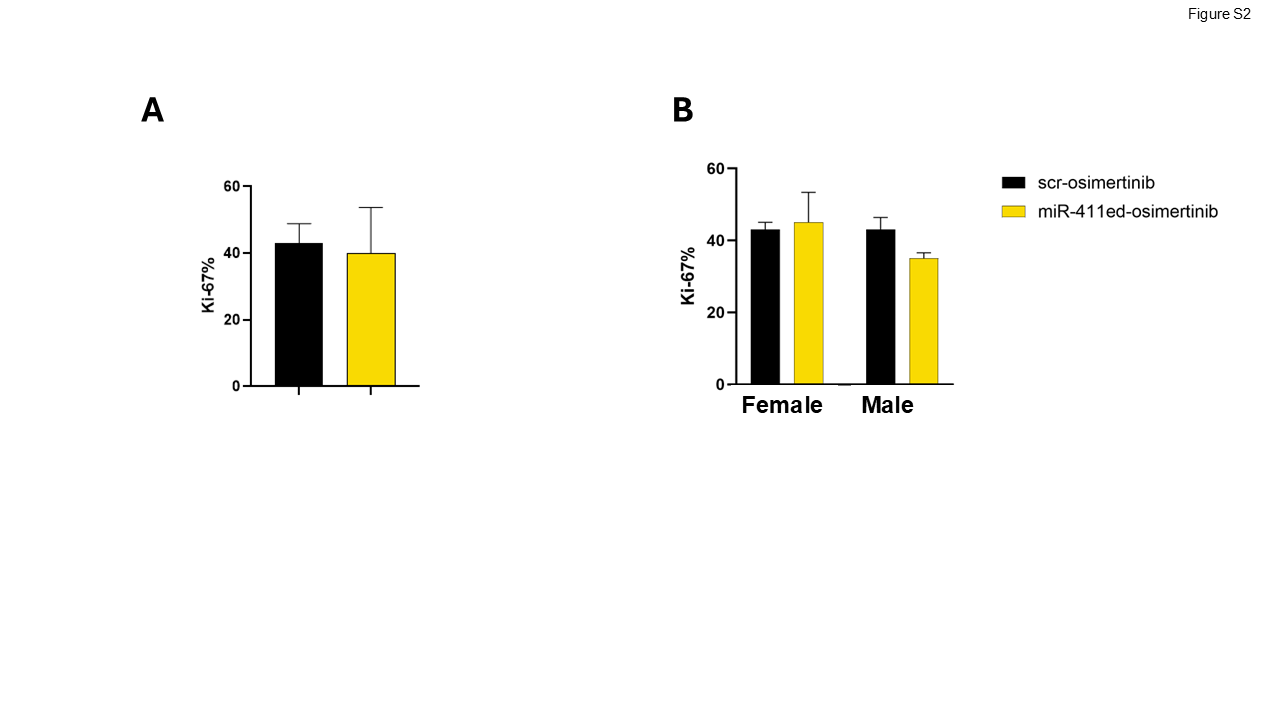
